# GOLIATH regulates LDLR availability and plasma LDL cholesterol levels

**DOI:** 10.1101/2020.12.09.415323

**Authors:** Bethan L. Clifford, Kelsey E. Jarrett, Joan Cheng, Angela Cheng, Marcus Seldin, Pauline Morand, Richard Lee, Angel Baldan, Thomas Q. de Aguiar Vallim, Elizabeth J. Tarling

**Affiliations:** Department of Medicine, Division of Cardiology, University of California Los Angeles, CA, USA; Department of Biological Chemistry, David Geffen School of Medicine at UCLA, University of California Los Angeles, CA, USA; Department of Biological Chemistry, University of California Irvine, CA, USA; Ionis Pharmaceuticals, Carlsbad, CA, USA; Edward A. Doisy Department of Biochemistry and Molecular Biology, Saint Louis University, St. Louis, MO, USA; Molecular Biology Institute, David Geffen School of Medicine at UCLA, University of California Los Angeles, CA, USA; Jonsson Comprehensive Cancer Center, David Geffen School of Medicine at UCLA, University of California Los Angeles, CA, USA

**Keywords:** E3 ligase, lipids, cholesterol, LDL receptor, ubiquitin

## Abstract

Increasing the availability of hepatic low-density lipoprotein receptors (LDLR) remains a major clinical target for reducing circulating plasma LDL cholesterol (LDL-C) levels. Here, we identify the molecular mechanism underlying genome-wide significant associations in the *GOLIATH* locus with plasma LDL-C levels. We demonstrate that GOLIATH is an E3 ubiquitin ligase that ubiquitinates the LDL Receptor resulting in redistribution away from the plasma membrane. Overexpression of GOLIATH decreases hepatic LDLR and increases plasma LDL-C levels. Silencing of *Goliath* using antisense oligonucleotides, germline deletion, or AAV-CRISPR *in vivo* strategies increases hepatic LDLR abundance and availability, thus decreasing plasma LDL-C. *In vitro* ubiquitination assays demonstrate RING-dependent regulation of LDLR abundance at the plasma membrane. Our studies identify GOLIATH as a novel post-translational regulator of LDL-C levels via modulation of LDLR availability, which is likely important for understanding the complex regulation of hepatic LDLR.

## Introduction

Elevated plasma low-density lipoprotein cholesterol (LDL-C) levels are a well-established risk factor for cardiovascular disease (1). Plasma LDL particle concentration is regulated by the balance between synthesis, from its very low-density lipoprotein (VLDL) precursor, and uptake from the circulation via the LDL receptor (LDLR) (2). The liver expresses over 70% of whole body LDLR and is the major determinant of plasma LDL-C concentrations (3, 4). As such, regulation of LDLR is under tight control by multiple complex pathways.

Human genetic studies have successfully identified loci containing genes previously implicated in regulating lipid levels, such as LDLR, HMGCR (3-Hydroxy-3-Methylglutaryl-CoA Reductase), and PCSK9 (Proprotein convertase subtilisin/kexin type 9). However, a significant number of additional loci have been associated with lipid levels that contain potential unrecognized regulators of lipid traits that have not been well characterized, and the causal genes remain to be identified. Further, the molecular mechanisms underlying such associations remain elusive. Understanding the molecular mechanisms driving these associations is key to leveraging loci identified through lipid genome-wide associations, which to date has been lacking.

GOLIATH is the human homolog of the *Drosophila* protease-associated (PA) domain-containing E3 ligase Goliath. E3 ligases are part of the cellular ubiquitin system that can not only control the selective degradation of proteins by the 26S proteasome, but can also modify proteins and regulate their cellular localization (11). E3 ligases catalyze transfer of ubiquitin to the target protein, thereby conferring specificity and regulation to these processes. Most E3 ligases are cytosolic, presumably to increase their exposure to substrates, E2 conjugating enzymes and ubiquitin, thus allowing for rapid regulation of target proteins. Despite there being over 600 RING E3 ligases (12), only a small number, that includes GOLIATH, contain a *bona fide* transmembrane domain (TMD) (13). The function and molecular targets of GOLIATH are unknown (14, 15).

GOLIATH, like the other members of the PA domain-containing E3 ligase sub-family, exhibits a distinct domain architecture, consisting of a signal peptide, a PA domain, a TMD, and a RING (Really Interesting New Gene) domain (16). Ubiquitination events mediated by *Drosophila* Goliath family members have been shown to regulate specific cellular events such as endocytosis (13, 17) and protein-protein interactions (18, 19). Given the highly conserved structural architecture between GOLIATH and other PA-TM-RING proteins, we hypothesized that GOLIATH may play a role in the endocytic recycling of membrane receptors that regulate plasma cholesterol, such as LDLR.

In the present study we identify GOLIATH as a novel post-translational regulator of hepatic LDLR and plasma LDL-C levels, thus explaining previously identified associations between the *GOLIATH* locus and plasma cholesterol levels in humans. Using a combination of molecular, biochemical, and *in vivo* metabolic analyses, we have determined that GOLIATH ubiquitinates the LDLR, reduces plasma membrane LDLR localization and increases plasma LDL-C levels. We also demonstrate, using three different independent *in vivo* approaches, that loss of GOLIATH results in increased hepatic levels of LDLR and reduced plasma LDL-C levels. We propose that this mechanism allows for the modulation of LDLR endocytosis and/or recycling, and therefore the regulation of available LDLR molecules at the cell surface.

## Results

### GOLIATH is associated with plasma LDL-C in both humans and mice

Human genome-wide association studies (GWAS) previously identified multiple single nucleotide polymorphisms (SNPs) associated with plasma LDL-C in intron 1 of *GOLIATH*, a locus not previously implicated in lipoprotein metabolism (9). Human and mouse GOLIATH protein are highly conserved and share 98% sequence identity suggesting conserved function between species. As such, we sought additional evidence for the involvement of GOLIATH in regulating LDL-C, by comparing the previous linkage results from human studies with results from a diverse panel of 120 inbred mouse strains known as the Hybrid Mouse Diversity Panel (HMDP; (20, 21). In this panel, we identified significant correlation between hepatic mRNA expression of *Goliath* and plasma lipid traits including total (Supplemental Fig. 1A), LDL (Supplemental Fig. 1B), and HDL (High-density lipoprotein) (Supplemental Fig. 1C) cholesterol levels. These data are consistent with other significant regulators of cholesterol, such as LDLR (*P*=2.1 × 10^−4^). Using a panel of tissues isolated from C57BL/6 mice we determined that *Goliath* is ubiquitously expressed in mice, with highest expression in metabolic tissues such as liver and adipose (Supplemental Fig. 1D).

**Figure 1.**
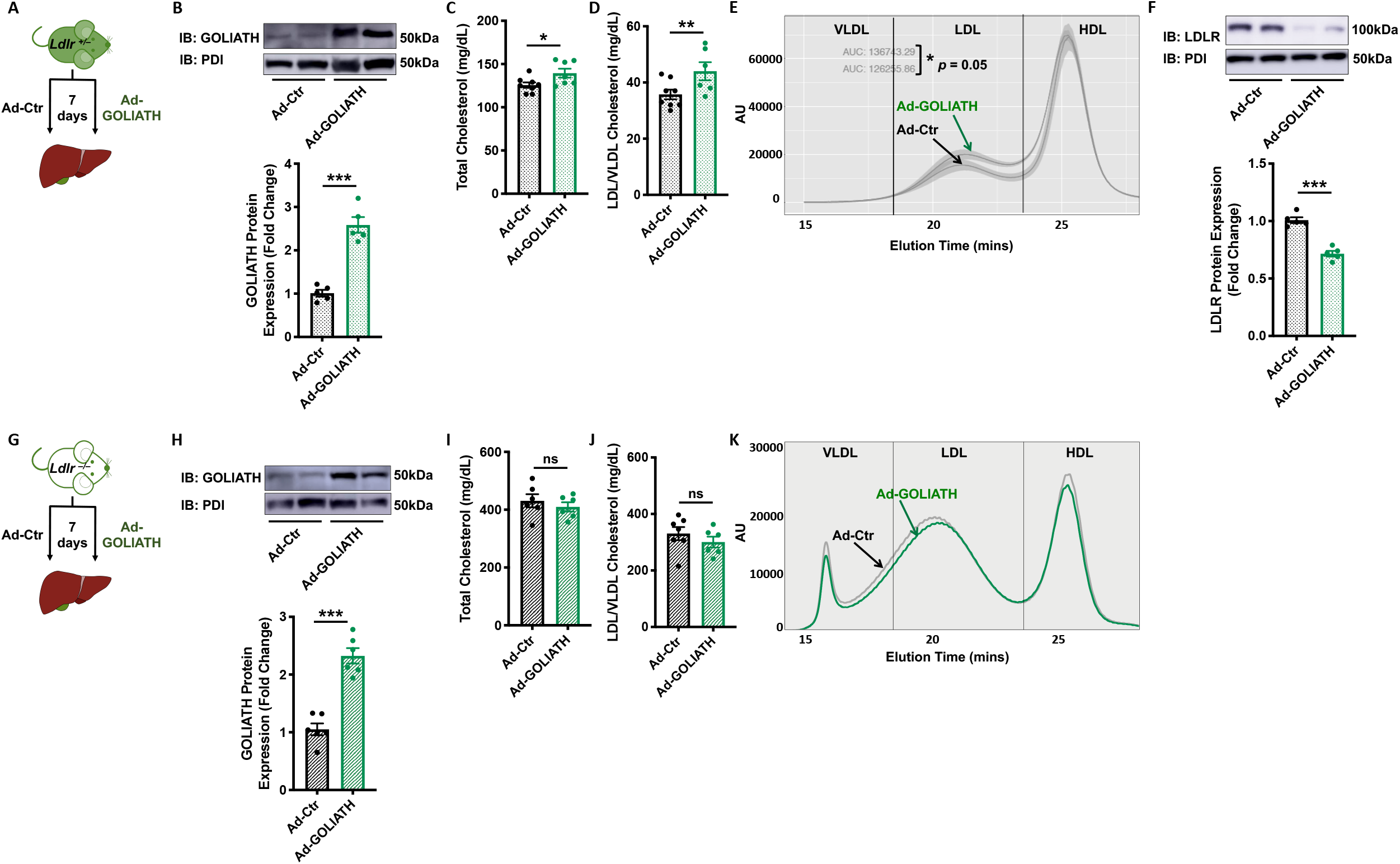
GOLIATH reduces plasma LDL-C levels *in vivo* in a process that is dependent on LDLR. **(A, G)** Experiment schematics. **(A)** *Ldlr*^*+/–*^ or **(G)** *Ldlr*^*–/–*^ mice were treated with control adenovirus (Ad-Ctr) or adenovirus overexpressing human GOLIATH (Ad-GOLIATH) for 7 days (*n*=5-8 mice/group). **(B, H)** Hepatic protein expression and accompanying densitometry of GOLIATH in **(B)** *Ldlr*^*+/–*^ or **(H)** *Ldlr*^*–/–*^ mice treated with Ad-Ctr or Ad-GOLIATH. Plasma total cholesterol **(C)**, LDL (LDL/VLDL) cholesterol **(D)**, and FPLC lipoprotein profiles **(E)** in *Ldlr*^*+/–*^ mice treated with Ad-Ctr or Ad-GOLIATH. **(F)** Hepatic LDLR protein expression and accompanying densitometry in *Ldlr*^*+/–*^ mice treated with Ad-Ctr or Ad-GOLIATH. Plasma total cholesterol **(I)**, LDL (LDL/VLDL) cholesterol **(J)**, and FPLC lipoprotein profiles **(K)** in *Ldlr*^*–/–*^ mice treated with Ad-Ctr or Ad-GOLIATH. Data are expressed as mean ±SEM with individual animals noted as dots. Statistical significance was determined by Student’s *t* test. * *p*<0.05, ** *p*<0.01, *** *p*<0.001. ns, not significant.

### GOLIATH reduces plasma LDL-C levels *in vivo* in a process that is dependent on LDLR

To determine if increases in expression of GOLIATH are associated with changes in plasma LDL-C levels, we generated adenovirus particles to overexpress human GOLIATH in the livers of mice. C57BL/6 mice are known to have low levels of circulating plasma LDL-C (22), so we first overexpressed human GOLIATH in LDLR heterozygous (*Ldlr*^*+/–*^) mice, a model with elevated circulating LDL-C levels owing to a 50% reduction in functional LDL receptors. Indeed, patients who are heterozygous for familial hypercholesterolemia (FH), and thus have only one functional *LDLR* gene, have substantially elevated plasma LDL-C levels (23, 24). *Ldlr*^*+/–*^ mice were injected with either control adenovirus (Ad-Ctr) or adenovirus expressing human *GOLIATH* (Ad-GOLIATH) for 7 days (Fig. 1A). GOLIATH overexpression (Fig. 1B) resulted in a significant increase in total (Fig. 1C) and non-HDL (LDL/VLDL) plasma cholesterol levels (Fig. 1D). No differences were seen in HDL-C or triglyceride levels (TAG, Supplemental Fig. 1E-F). Fast performance liquid chromatography (FPLC) analysis of plasma samples from individual mice demonstrated a significant increase in the LDL-C fraction (Fig. 1E). Elevated circulating levels of LDL-C can arise due to reduced or defective hepatic LDLR levels and/or activity. Therefore, we quantified hepatic LDLR mRNA and protein levels. Overexpression of GOLIATH protein did not affect hepatic *Ldlr* mRNA levels (Supplemental Fig. 1G), but significantly decreased hepatic LDLR protein levels (Fig. 1F). To determine if the GOLIATH-mediated increase in LDL-C is dependent on the presence of hepatic LDLR, we overexpressed GOLIATH in LDLR knockout (*Ldlr*^*–/–*^) mice (Fig. 1G). Overexpression of GOLIATH (Fig. 1H) in *Ldlr*^*–/–*^ mice did not change plasma total cholesterol (Fig. 1I, 1K), LDL/VLDL-C (Fig. 1J-K), HDL-C, or TAG levels (Supplemental Fig. 1H-I), demonstrating that LDLR is required for the GOLIATH-dependent increase in LDL-C levels.

### Efficacy of GOLIATH-mediated regulation of plasma cholesterol levels is dependent on hepatic LDLR abundance

Based on these data, we hypothesized that overexpression of GOLIATH may have more potent effects on plasma LDL-C in mouse models that exhibit elevated endogenous levels of hepatic LDLR protein compared to *Ldlr*^*+/–*^ mice. First, we overexpressed GOLIATH in wildtype mice that have fully intact and functioning LDL receptors (Fig. 2A). GOLIATH overexpression (Fig. 2B) resulted in significantly increased total (Fig. 2C) and non-HDL (LDL/VLDL) plasma cholesterol levels (Fig. 2D). Plasma HDL-C and triglyceride levels were either unchanged (Supplemental Fig. 2A) or modestly increased (Supplemental Fig. 2B). FPLC analysis of plasma samples from individual mice demonstrated a significant increase in the LDL-C fraction (Fig. 2E). Consistent with what we observed in *Ldlr*^*+/–*^ mice (Fig. 1F), GOLIATH overexpression also significantly reduced hepatic LDLR protein levels (Fig. 2F).

**Figure 2.**
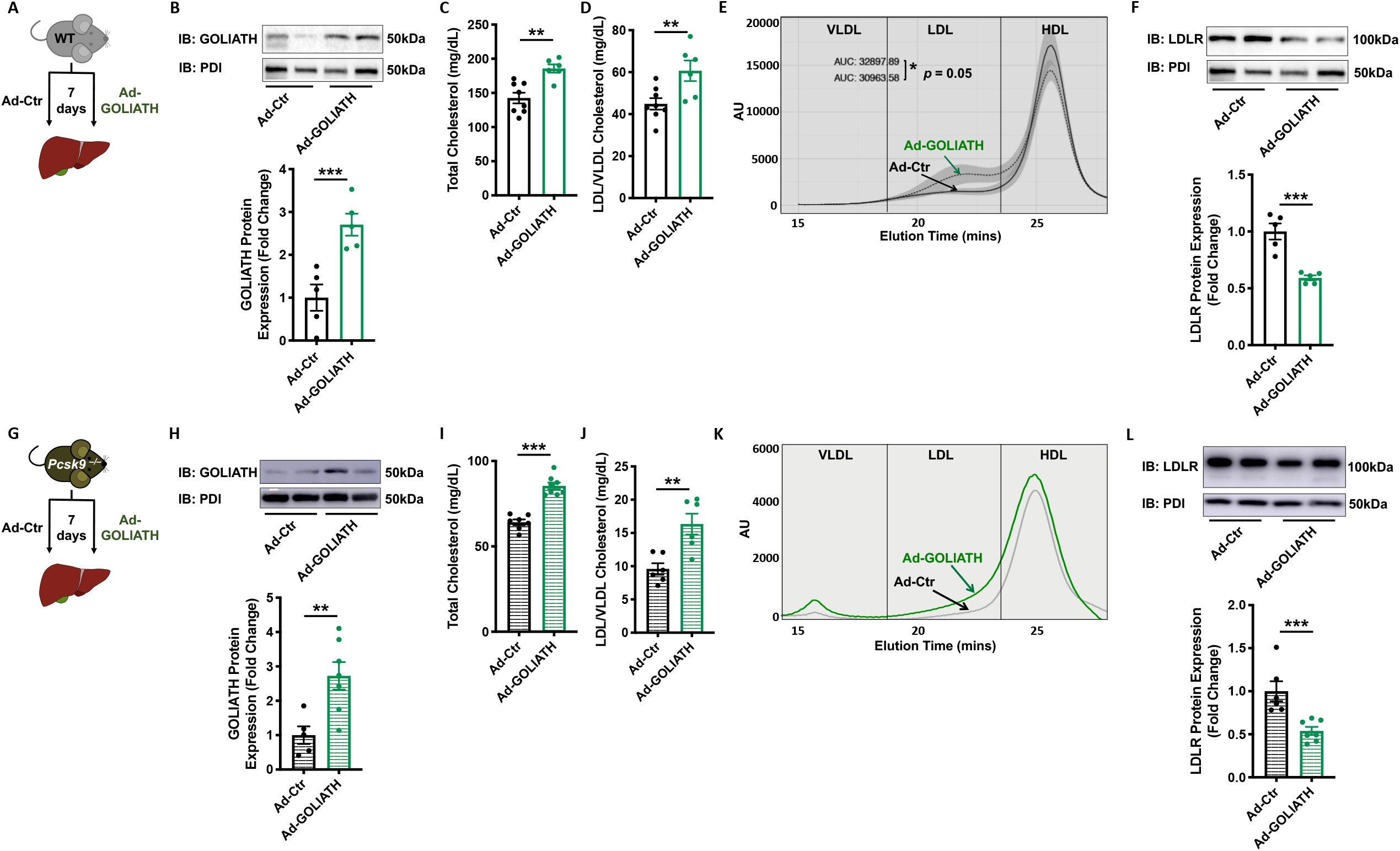
Efficacy of GOLIATH-mediated regulation of plasma cholesterol levels is dependent on hepatic LDLR abundance. **(A, G)** Experiment schematics. **(A)** Wildtype (WT) or **(G)** *Pcsk9*^*–/–*^ mice were treated with control adenovirus (Ad-Ctr) or adenovirus overexpressing human GOLIATH (Ad-GOLIATH) for 7 days (*n*=5-8 mice/group). **(B, H)** Hepatic protein expression and accompanying densitometry of GOLIATH in **(B)**wildtype or **(H)** *Pcsk9*^*–/–*^ mice treated with Ad-Ctr or Ad-GOLIATH. Plasma total cholesterol **(C)**, LDL (LDL/VLDL) cholesterol **(D)**, and FPLC lipoprotein profiles **(E)** in wildtype mice treated with Ad-Ctr or Ad-GOLIATH. **(F, L)** Hepatic LDLR protein expression in wildtype mice **(F)** and *Pcsk9*^*–/–*^ mice **(L)** treated with Ad-Ctr or Ad-GOLIATH. Plasma total cholesterol **(I)**, LDL (LDL/VLDL) cholesterol **(J)**, and FPLC lipoprotein profiles **(K)** in *Pcsk9*^*–/–*^ mice treated with Ad-Ctr or Ad-GOLIATH. Data are expressed as mean ±SEM with individual animals noted as dots. Statistical significance was determined by Student’s *t* test. * *p*<0.05, ** *p*<0.01, *** *p*<0.001. ns, not significant.

Next, we overexpressed GOLIATH in *Pcsk9*^*–/–*^ mice, a model of increased hepatic LDL receptors (25, 26). PCSK9 normally binds to the LDLR/LDL-C complex as it is being internalized and signals for the LDLR to be degraded in the lysosome, instead of being recycled back to the plasma membrane (27). Consequently, compared to *Pcsk9*^*+/+*^ mice, *Pcsk9*^*–/–*^ mice have elevated levels of hepatic LDLR protein and significantly reduced levels of plasma LDL-C (26).

Overexpression of GOLIATH in *Pcsk9*^*–/–*^ mice (Fig. 2G-H) resulted in significantly decreased LDLR protein levels (Fig. 2L) and increased plasma total cholesterol (Fig. 2I, 2K), LDL/VLDL-C (Fig. 2J-K), and HDL-C levels (Supplemental Fig. 2C), with no change in plasma triglyceride levels (Supplemental Fig. 2D). Together these data show that overexpression of GOLIATH in *Ldlr*^*+/–*^, WT or *Pcsk9*^*–/–*^ mice, increased LDL/VLDL-C levels by 25%, 36% and 70%, respectively, (Fig. 1D-E, Fig. 2D-E, 2J-K), suggesting that the efficacy of GOLIATH for increasing LDL/VLDL-C is enhanced when basal hepatic LDLR levels are elevated.

Overexpression of GOLIATH did not affect levels of the endocytosed receptors EGFR, LRP-1, and Transferrin receptor (Supplemental Fig. 2E), suggesting specificity for LDLR rather than gross changes in endocytosis. Taken together, these data suggest that overexpression of GOLIATH increases plasma LDL-C by a process dependent on the expression and availability of hepatic LDLR.

### GOLIATH is a RING-dependent E3 ligase that ubiquitinates LDLR and redistributes LDLR away from the plasma membrane

Having demonstrated that plasma LDL-C and LDLR levels are modulated by GOLIATH *in vivo*, we next sought to determine the molecular mechanism underlying this regulation. Given that GOLIATH is reported to function as an E3 ubiquitin ligase (14), we hypothesized that LDLR may be ubiquitinated by GOLIATH. To test this hypothesis, we transfected HEK293 cells with plasmids encoding GFP-tagged LDLR, HA-tagged ubiquitin, and, where indicated, an N-terminally FLAG-tagged human GOLIATH. Since the coding sequence for GOLIATH contains a predicted signal peptide (amino acids 1-27), and addition of a tag on the carboxy-terminal end of the protein renders it non-functional (data not shown), we engineered a 3xFLAG-tag immediately after the signal peptide. As shown in Fig. 3A, total cellular LDLR protein was decreased in cells co-expressing GOLIATH (Fig. 3A; Input) which was accompanied by greatly increased ubiquitination of LDLR (Fig. 3A; Ubiquitination).

**Figure 3.**
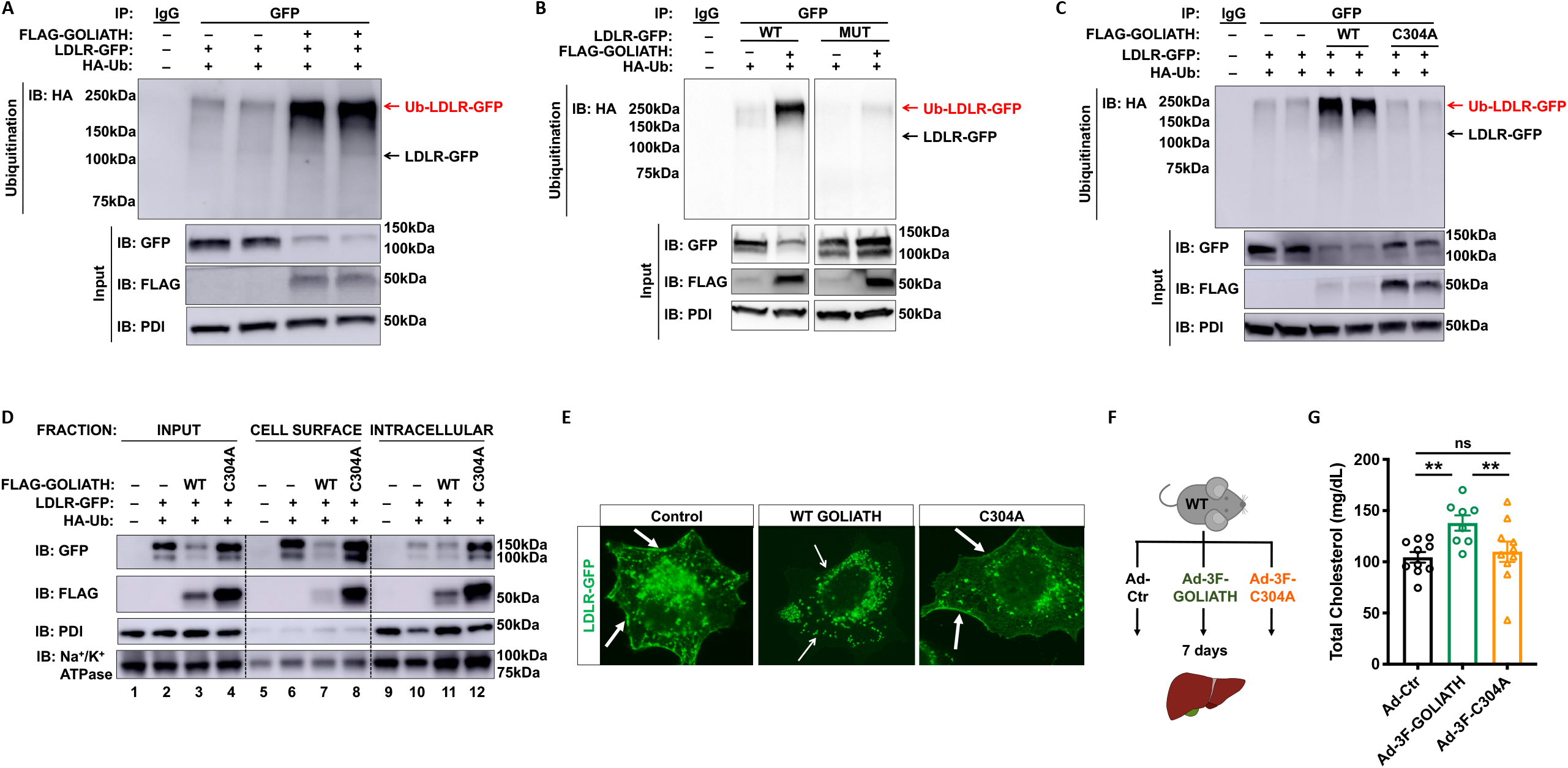
GOLIATH is a RING-dependent E3 ligase that ubiquitinates LDLR and redistributes LDLR away from the plasma membrane. **(A)** HEK293 cells were co-transfected with LDLR-GFP, FLAG-GOLIATH, and HA-Ubiquitin expression plasmids. After 36 hours, lysates were subjected to immunoprecipitation (IP) with anti-GFP and immunoblotting (IB) with anti-HA to detect ubiquitinated proteins (upper panel). Total cell lysates (input; lower panels) were processed for western blotting with antibodies to anti-GFP, anti-FLAG, or anti-PDI. GFP, green fluorescent protein; HA, hemagglutinin; PDI, protein disulfide isomerase; Ub, ubiquitin. **(B)** HEK293 cells were co-transfected with wildtype (WT) or mutant (MUT) LDLR-GFP (K811/816/830RC839A), FLAG-GOLIATH, and HA-Ubiquitin. After 36 hours, lysates were subjected to IP and IB as in (A). **(C)** HEK293 cells were co-transfected with LDLR-GFP, wildtype (WT) FLAG-GOLIATH or mutant FLAG-GOLIATH (C304A) and HA-Ubiquitin expression plasmids. After 36 hours, lysates were subjected to IP and IB as in (A). **(D)** HEK293 cells were co-transfected with LDLR-GFP, wildtype (WT) FLAG-GOLIATH or mutant FLAG-GOLIATH (C304A) and HA-Ubiquitin expression plasmids. After 36 hours, cells were incubated with biotin at 4°C to label cell surface proteins. Total cell lysates (input; lanes 1-4), biotinylated proteins (membrane; lanes 5-8), and unmodified proteins (intracellular; lanes 9-12) were processed for western blotting with antibodies to anti-GFP, anti-FLAG, anti-PDI (intracellular) and anti-Na^+^/K^+^ ATPase (plasma membrane). **(E)** Immunofluorescence images of HEK293 cells co-transfected with LDLR-GFP and either wildtype (WT) FLAG-GOLIATH or mutant FLAG-GOLIATH (C304A) Original magnification 63X. White arrows indicate plasma membrane. **(F)** Experiment schematic. Wildtype (WT) mice were treated with control adenovirus (Ad-Ctr), or adenovirus overexpressing human GOLIATH with an N-terminal 3xFLAG epitope tag (Ad-3F-GOLIATH) or 3F-GOLIATH with a single point mutation in the RING domain (Ad-3F-C304A) for 7 days (*n*=8-10 mice/group). **(G)** Plasma total cholesterol in mice treated as in (f). Data are expressed as mean ±SEM with individual animals noted as dots. Statistical significance was determined by one-way ANOVA. ** *p* < 0.01, *** *p* < 0.001, ns. Not significant.

LDLR has multiple highly conserved potential ubiquitination sites within its cytoplasmic tail (Lys^811^, Lys^816^, Lys^830^, and Cys^839^). We repeated the experiments described above but using an LDLR construct where these four residues were mutated to arginine or alanine, respectively. Combined mutation of all four residues (K811/816/830R/C839A) prevented both ubiquitination of LDLR and the loss of LDLR-GFP protein (Fig. 3B). Consistent with *in vivo* observations, overexpression of GOLIATH had no effect on levels of additional endocytosed receptors EGFR, TfR, and the related lipoprotein receptor LRP-1 (Supplemental Fig. 3A), further suggesting that this ubiquitination event is specific to LDLR and not a gross change in receptors known to be internalized via clathrin-mediated endocytosis.

Having determined that GOLIATH requires the cytoplasmic tail of LDLR to be intact for a ubiquitination event, we next turned to assessing whether the E3 ubiquitin ligase function of GOLIATH was essential for this observation. The catalytic domain required for ubiquitination activity of RING E3 ligases is the RING domain and specific mutations in this domain have been shown to impair ubiquitination of target proteins (12, 28). Consistent with this observation, GOLIATH containing a single point mutation in the RING domain (Cys^304^ → Ala^304^) failed to ubiquitinate LDLR or lower LDLR protein levels (Fig. 3C). Further, the levels of FLAG-tagged GOLIATH were elevated when the RING domain was mutated, compared to wildtype GOLIATH (Fig. 3C), suggesting that GOLIATH can also catalyze its own ubiquitination and degradation. Such self-ubiquitination is in agreement with previously published reports of GOLIATH (14, 15) and other RING E3 ligases.

A significant percent of LDLR protein is normally expressed at the cell surface which is important for its function in the endocytic pathway that internalizes LDL-C particles (29). To determine whether GOLIATH affects LDLR abundance at the cell surface, we transfected HEK293 cells with plasmids encoding GFP-tagged LDLR, HA-tagged ubiquitin, and N-terminally FLAG-tagged WT or mutant (C304A) GOLIATH. Transfected cells were exposed to biotin to label cell surface proteins before quenching the reaction to prevent subsequent modification of intracellular proteins released after cell lysis. As expected, WT FLAG-GOLIATH but not mutant FLAG-GOLIATH^C304A^, reduced GFP-tagged LDLR protein levels in whole cell lysates (Fig. 3D; IB:GFP lanes 2-4). WT GOLIATH had a marked effect on the abundance of cell surface levels of GFP-tagged LDLR (Fig. 3D; IB:GFP lane 7 *vs*. 6). In contrast, cell surface levels of GFP-LDLR actually increased slightly in cells co-expressing following expression of mutant GOLIATH^C304A^ (Fig. 3D; IB:GFP lane 8 *vs*. 6). Analysis of intracellular LDLR levels indicated that there was no significant change in intracellular LDLR levels following co-expression of WT GOLIATH (Fig. 3D; IB:GFP lane 11 *vs*. 10), but an increase following expression of mutant GOLIATH^C304A^ (Fig. 3D; IB:GFP lane 12 *vs*. 10). The finding that intracellular levels of LDLR increased following co-expression of mutant GOLIATH^C304A^ suggests that the normal degradation of the LDLR might be impaired under these latter conditions. Together, these data suggest GOLIATH had proportionally greater effects on LDLR abundance at the cell surface.

In addition to signaling for degradation in the proteasome, protein ubiquitination can also serve as an internalization signal for specific proteins at the plasma membrane (13, 30). As effects of GOLIATH on LDLR protein levels were greater at the cell surface, we hypothesized that the ubiquitination of LDLR by GOLIATH may result in redistribution of LDLR away from the cell surface, in addition to possibly increasing degradation of LDLR. Immunohistochemical staining showed that co-expression of GOLIATH dramatically decreased the levels of LDLR protein at the cell surface (Fig. 3E; middle panel). Although there was still LDLR protein in intracellular compartments (Fig. 3E; middle panel), quantification of mean fluorescent intensity demonstrated a significant reduction in GFP fluorescence indicating that total levels of GFP-tagged LDLR were decreased with co-expression of GOLIATH (Supplemental Fig. 3B). In contrast to WT GOLIATH, cells expressing mutant GOLIATH^C304A^ continued to express high levels of LDLR both at the cell surface and in intracellular compartments (Fig. 3E; right panel and Supplemental Fig. 3B).

Finally, to assess whether the E3 ubiquitin ligase function of GOLIATH was required for its ability to increase plasma LDL-C *in vivo*, we treated WT mice with either control or adenovirus overexpressing FLAG-tagged WT GOLIATH or GOLIATH containing the C304A point mutation in the RING domain (Fig. 3F). Consistent with previous studies (Fig. 1-2), FLAG-GOLIATH overexpression in WT mice (Fig. 3F, Supplemental Fig. 3D-E) increased total and LDL cholesterol levels (Fig. 3G, Supplemental Fig. 3C) and this was associated with a decrease in LDLR protein (Supplemental Fig. 3D, 3F). In contrast, expression of mutant GOLIATH^C304A^ (Supplemental Fig. 3D-E) failed to alter plasma cholesterol levels (Fig. 3G) or LDLR protein (Supplemental Fig. 3D, 3F). Collectively, these experiments demonstrate that GOLIATH regulates LDL-C through its function as a ubiquitin ligase, and that the consequence of this ubiquitination is likely a combination of redistribution of LDLR from the plasma membrane and degradation of LDLR protein.

### Partial knockout of *Goliath* in mice results in increased hepatic LDLR and reduced plasma cholesterol

Thus far, we have shown that GOLIATH gain-of-function results in increased plasma LDL-C levels. To carry out the reverse loss-of-function, we first generated mice with a targeted disruption of *Goliath*. Germline transmission was confirmed by PCR genotyping (Supplemental Fig. 4A) for a minimum of ten litters, as previously described (31). When heterozygous *Goliath*^*+/–*^ mice were crossed with one another, pups were obtained at the ratio of 1 *Goliath*^*+/+*^: 2 *Goliath*^*+/–*^: 0.6 *Goliath*^*–/–*^ (n=35 litters, 204 pups total, mean litter size 5.8 pups per litter) which differs from the expected Mendelian ratio of 1:2:1 (Fig. 4A). These data suggest that homozygous deletion of *Goliath* is partially embryonic lethal.

**Figure 4.**
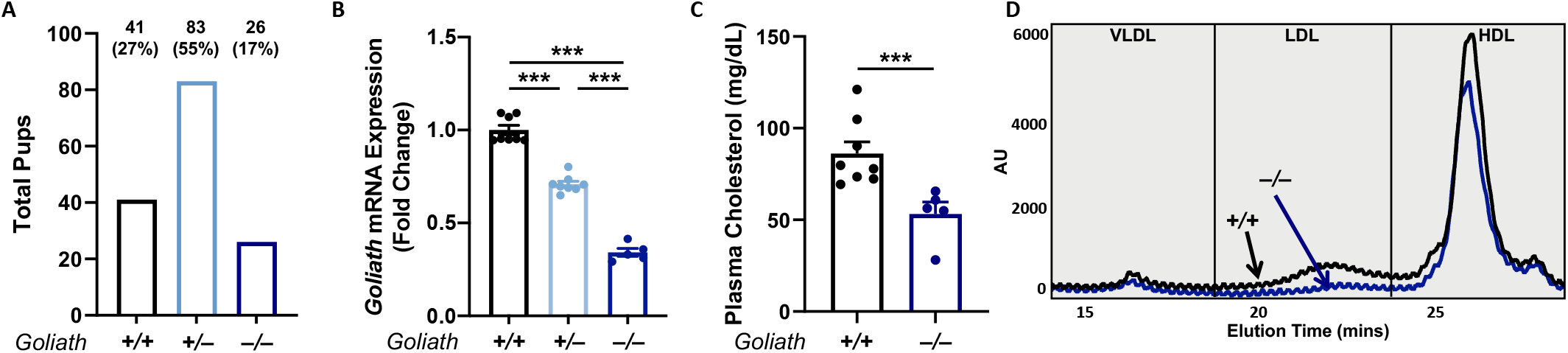
Knockout of *Goliath* results in decreased plasma LDL-C. **(A)** Total number of pups obtained for each genotype from heterozygous matings showing that homozygous nulls (–/–) are obtained at less than the expected mendelian ratio (n=35 litters, 204 pups total, mean litter size 5.8). **(B)** Hepatic mRNA expression of *Goliath*, **(C)** total plasma cholesterol, and **(D)**FPLC lipoprotein profiles of 10-week old *Goliath*^*+/+*^ and *Goliath*^*–/–*^ mice. (n=5-8 mice/genotype). Data are expressed as mean ±SEM with individual animals represented as dots. Significance was measured by one-way ANOVA. * *p* < 0.05, *** *p* < 0.001. ns, not significant.

We next investigated the phenotype of the *Goliath*^*–/–*^ mice that survived birth. Surviving *Goliath*^*–/–*^ pups were able to nurse, feed, and mature into adults, and the body weight of adult mice at 10 weeks of age was not significantly different from either heterozygous (*Goliath*^*+/–*^) or wildtype (*Goliath*^*+/+*^) mice (Supplemental Fig. 4B). Gene expression analysis of livers from *Goliath*^*+/+*^, *Goliath*^*+/–*^, and *Goliath*^*–/–*^ mice demonstrated that *Goliath*^*–/–*^ mice had 30% residual *Goliath* mRNA expression (Fig. 4B), suggesting that complete deletion of *Goliath* is not compatible with survival, and surviving mice have escaped entire loss of *Goliath*. Similarly, *Goliath*^*+/–*^ mice only experienced a 30% knockdown of mRNA expression, significantly lower than the 50% that would be expected of heterozygous animals (Fig. 4B). Importantly, plasma lipid analysis in *Goliath*^*–/–*^ mice showed a significant reduction in circulating plasma total cholesterol (Fig. 4C) and these changes appeared to be present in the LDL fraction (Fig. 4D). Consistent with such a decrease in plasma LDL-C, there was a concomitant increase in LDLR protein expression in *Goliath*^*–/–*^ mice compared to their wild-type littermates (Supplemental Fig. 4C).

### Liver-specific disruption of *Goliath* reduces plasma cholesterol levels

Given that embryonic deletion of *Goliath* is not compatible with survival and 70% of whole body LDLR is expressed in the liver, we next sought to determine whether acute reductions in GOLIATH in the livers of adult mice reduced plasma LDL-C levels. We used an all-in-one AAV-CRISPR vector expressing a small guide RNA (gRNA) targeting exon 5 of *Goliath*, and Cas9 from *Staphylococcus aureus* (SaCas9) (Supplemental Fig. 5A). We targeted exon 5 because exon 5 encodes the RING domain, which is critical for GOLIATH function (Fig. 3). The CRISPR system was packaged into AAV 2/8 which has a high tropism for liver (35, 36), and we increased liver-specificity further by expressing SaCas9 under the control of a liver-specific promoter (Supplemental Fig. 5A). C57BL/6 mice were assigned into groups and blood samples drawn to ensure equivalent levels of plasma cholesterol at baseline (Supplemental Fig. 5B). Mice were injected with either control AAV-CRISPR or *Goliath* AAV-CRISPR (Fig. 5A) at a dose of 5×10^11^ genome copies per animal. Two-week treatment with either Control or *Goliath* AAV-CRISPR had no effect on body weight (Supplemental Fig. 5C). Hepatic *Goliath* expression was decreased by 70% in mice treated with *Goliath* AAV-CRISPR (Fig. 5B). To ensure comparable viral load had been achieved in both groups, we performed PCR to amplify within SaCas9 (Supplemental Fig. 5D) and measured viral genome copies by qPCR (Supplemental Fig. 5E). To confirm the decreases seen in *Goliath* mRNA were the consequence of CRISPR-mediated gene disruption, we sequenced samples from control- and *Goliath*-AAV-CRISPR treated animals and performed editing analyses using Synthego ICE. As expected, we detected INDELs in *Goliath* exon 5 in mice treated with *Goliath* AAV-CRISPR (Supplemental Fig. 5F). These analyses also show increased values for discordant sequences in these mice when compared to control treated animals (Supplemental Fig. 5G), suggesting some INDELs are affecting an area larger than the 8bp measurement window. Together, these data indicate that a proportion of the remaining mRNA is disrupted, and that the observed decrease in *Goliath* mRNA levels are the consequence of CRISPR gene editing.

**Figure 5.**
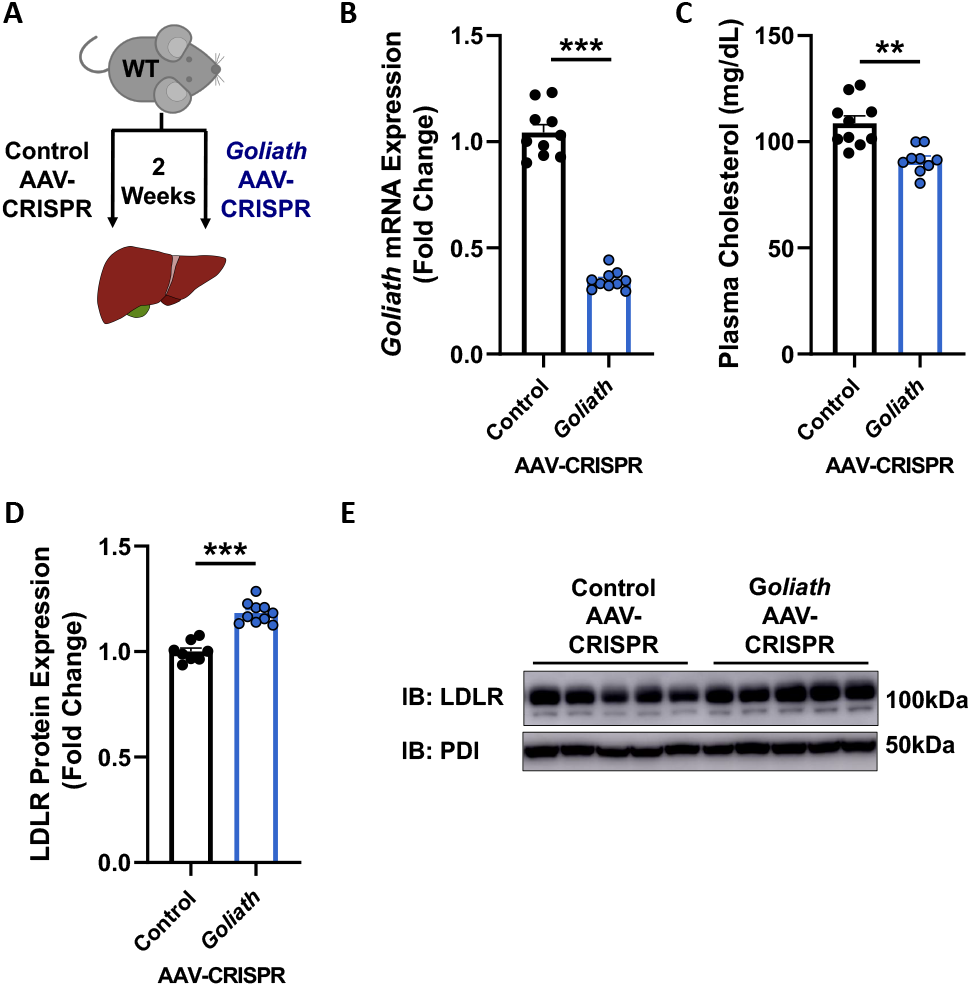
Liver-specific disruption of *Goliath* reduces plasma cholesterol levels. **(A)** C57BL/6 wildtype (WT) mice were treated with 5 × 10^11^ genome copies of Control AAV-CRISPR or *Goliath* AAV-CRISPR for 2 weeks (n=8-10/group). **(B)** Hepatic *Goliath* mRNA expression in wildtype mice treated as in (A). **(C)** Plasma total cholesterol in wildtype mice treated as in (A). **(D, E)** Hepatic LDLR protein expression and accompanying densitometry of mice treated as in (A). Data are expressed as mean ±SEM with individual animals represented as dots. Significance was measured by students t-test. ** *p*<0.01, *** *p*<0.001.

The AAV-CRISPR-mediated decrease in *Goliath* mRNA expression was accompanied by significantly decreased plasma cholesterol levels (Fig. 5C). Consistent with previous data in surviving *Goliath*^*–/–*^ mice (Fig. 4), we show increased LDLR protein levels in mice with reduced hepatic *Goliath* expression (Fig. 5D-E). Taken together, our data show that acute disruption of hepatic *Goliath* in adult mice results in increased LDLR and decreased plasma LDL-C levels.

### Silencing endogenous *Goliath* increases LDLR protein levels and decreases plasma LDL-C levels

The studies described above demonstrate that reducing hepatic *Goliath* mRNA expression by at least 70% either by gene targeting (Fig. 4) or CRISPR/Cas9 (Fig. 5) is sufficient to increase LDLR protein levels and lower plasma LDL-C levels. To assess the translational potential of these studies we utilized an antisense oligonucleotide (ASO; Ionis Pharmaceuticals) to silence *Goliath* for four weeks (Fig. 6A). In contrast to other genetic approaches to disrupt *Goliath* (Fig. 4-5), treatment with *Goliath* ASO for four weeks reduced hepatic *Goliath* mRNA levels by more than 90% (Fig. 6B). *Goliath* silencing caused significant decreases in total (Fig. 6C) and LDL (Fig. 6D-E) cholesterol. These decreases in plasma LDL-C concentrations were accompanied by significant increases in hepatic LDLR protein levels (Fig. 6F). These data demonstrate that silencing *Goliath* in adult mice using an ASO compound results in increased LDLR protein levels and lower circulating plasma LDL-C levels. Next, to determine whether these effects on LDL-C were dependent on the availability of LDLR, we repeated our ASO silencing experiment in *Ldlr*^*–/–*^ mice (Fig. 6G). As observed in wildtype mice treated with *Goliath* ASO (Fig. 6B), *Goliath* mRNA expression in ASO treated *Ldlr*^*–/–*^ mice was decreased by nearly 90% (Fig. 6H). However, silencing *Goliath* in *Ldlr*^*–/–*^ mice had no effect on plasma total and LDL-C concentrations (Fig. 6I-J), confirming that the effects on plasma cholesterol observed with silencing *Goliath* require expression of LDLR.

**Figure 6.**
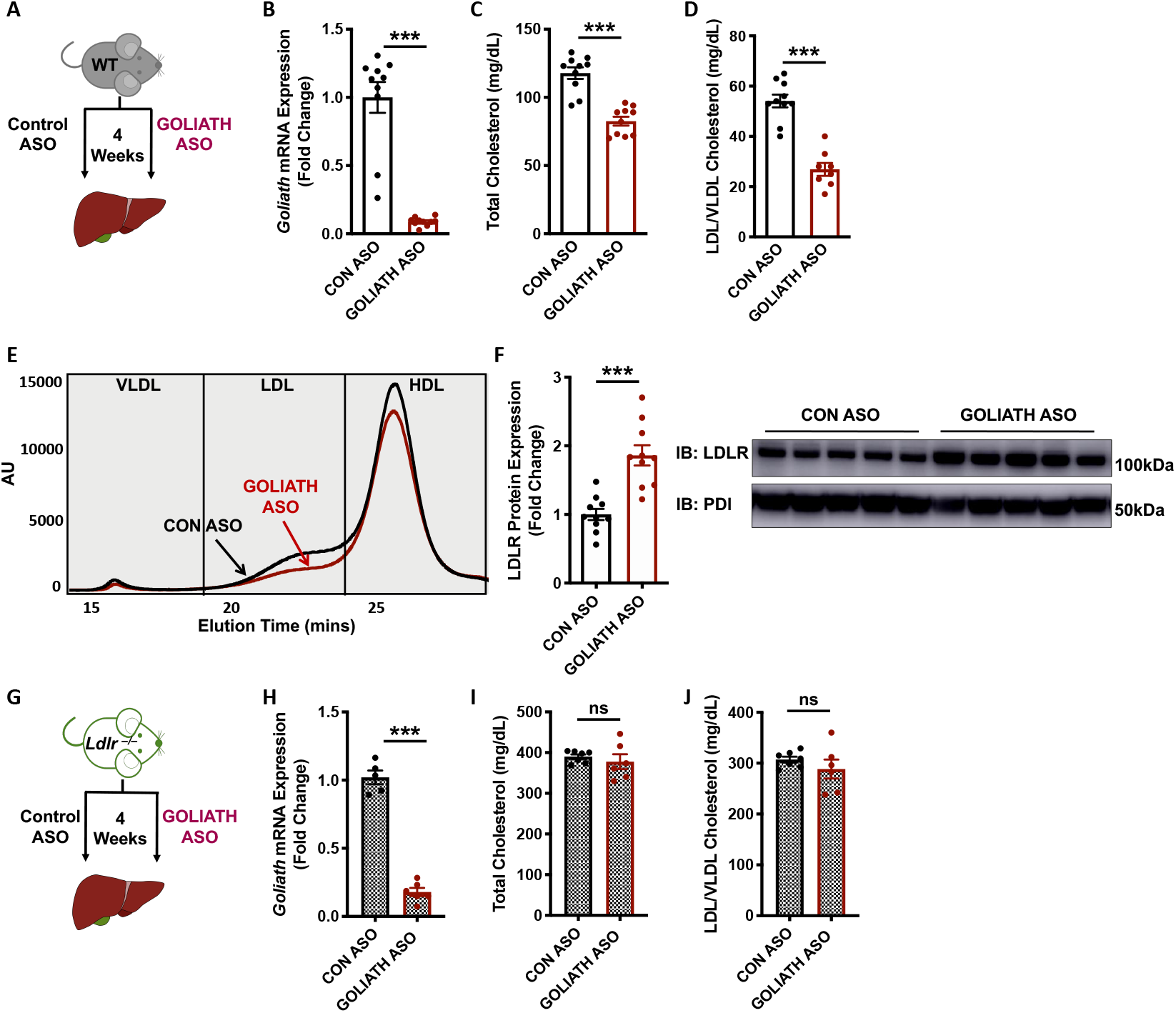
Antisense oligonucleotide treatment to silence hepatic *Goliath* increases hepatic LDLR and decreases plasma cholesterol levels. **(A)** C57BL/6 wildtype (WT) mice were treated with GOLIATH antisense oligonucleotide (ASO) for 4 weeks (*n*=10 mice/group). **(B)** Hepatic mRNA expression of *Goliath* in wildtype mice treated as in (A). **(C-E)** Plasma total cholesterol **(C)**, LDL (LDL/VLDL) cholesterol **(D)**, and FPLC lipoprotein profiles **(E)** in wildtype mice treated as in (A). **(F)** Hepatic LDLR protein expression in wildtype mice treated as in (A). **(G)** *Ldlr*^*–/–*^ mice were treated with GOLIATH ASO for 4 weeks. **(H)** Hepatic *Goliath* mRNA expression in *Ldlr–/–* mice treated as in (G). **(I-J)** Plasma total cholesterol **(I)** and LDL (LDL/VLDL) cholesterol **(J)** in *Ldlr–/–* mice treated as in (G). Data are expressed as mean ±SEM with individual animals noted as dots. Significance was measured by Student’s *t*-test. ** *p*<0.01, *** *p*<0.001, ns, not significant.

## Discussion

Genome-wide association studies are a powerful tool for identifying novel regulatory loci for many diseases. However, in most cases the molecular link explaining the association between the predicted loci and the disease remains unknown. Prior human genetic data suggested that GOLIATH plays a role in regulating plasma LDL-C levels as multiple SNPs in the *GOLIATH* locus strongly associate with LDL-C levels in humans (9). In the current study, we have identified the elusive mechanism underlying these previously observed associations and provide data identifying the molecular mechanism by which GOLIATH regulates plasma LDL-C. We utilize mice with gain-of function of GOLIATH, together with *in vitro* studies to demonstrate that GOLIATH ubiquitinates and redistributes LDLR away from the plasma membrane resulting in decreased levels of cell surface LDLR. We hypothesize that as a result LDLR-mediated clearance of LDL from the plasma is impaired during GOLIATH overexpression leading to elevated plasma LDL-C levels. Using three separate approaches to reduce *Goliath* expression *in vivo*, we further provide data demonstrating that GOLIATH loss-of-function leads to increased LDLR abundance and reduced plasma LDL-C levels.

GOLIATH is a member of the RING (Really Interesting New Gene) domain family of E3 ubiquitin ligases, which promote the transfer of ubiquitin from ubiquitin-conjugating enzymes (E2s) to lysine residues in target proteins (12). In the present study we show that GOLIATH ubiquitinates the LDLR and redistributes LDLR protein away from the plasma membrane. This ubiquitination of LDLR was dependent on both the catalytic RING domain of GOLIATH, and specific residues in the cytoplasmic tail of LDLR.

Earlier studies demonstrated that IDOL, another RING E3 ubiquitin ligase, also ubiquitinates LDLR (37). Importantly, there are a number of contrasting features between GOLIATH and IDOL. GOLIATH, unlike IDOL, is a member of the unique PA-TM-RING family of RING E3 ubiquitin ligases that contain a transmembrane (TM) domain. Previous studies of other PA-TM-RING ligases RNF13 and RNF167 have shown that these ligases are localized to both the plasma membrane and endosomes (13) and that this localization dictates target specificity and facilitates endocytic recycling of target proteins. IDOL, however, lacks a TM domain, is cytoplasmic, and contains a FERM binding domain critical for its interaction with LDLR family members. Finally, in contrast to GOLIATH, deletion of IDOL in mice had no effect on hepatic LDLR protein or plasma LDL-C levels, having more pronounced effects in peripheral tissues (38). Based on data presented in the current report, we postulate that GOLIATH functions by ubiquitinating LDLR at the plasma membrane to signal for receptor internalization, and/or by ubiquitinating endosomal LDLR to prevent recycling to the cell surface. These differences suggest that GOLIATH and IDOL may have important, contrasting roles in the regulation of hepatic LDLR and plasma LDL-C.

PCSK9 is an important post-translational regulator of LDLR levels (26, 39). Gain- and loss-of function mutations in PCSK9 have been identified in humans and PCSK9 monoclonal antibodies have been approved for the treatment of elevated plasma LDL-C levels (6, 40). The ability of GOLIATH to regulate LDLR and plasma LDL-C levels was preserved and enhanced in *Pcsk9*^*–/–*^ mice that have elevated LDLR protein levels, suggesting that GOLIATH’s effects on LDLR protein are independent of PCSK9. PCSK9 is known to have a complex reciprocal regulatory relationship with LDLR (25, 41). Studies have shown that PCSK9 is most effective in targeting the LDLR when the LDLR is bound to LDL particles (42, 43), and this may require the presence and/or availability of ApoE (44). Alternative means of post-translationally increasing the availability of unbound LDLR, such as through GOLIATH, are likely to further promote/enhance clearance of LDL-C and are therefore attractive additional therapeutic targets. ASO silencing of *Goliath* resulted in a 50% decrease in LDL-C, a change comparable to inhibiting PCSK9, further supporting that GOLIATH is a significant regulator of LDLR and LDL-C.

Whole body disruption of *Goliath* expression was incomplete and surviving pups exhibited only a 70% decrease in *Goliath* mRNA, suggesting that some residual expression of *Goliath* is necessary for survival. This is in agreement with publicly available human data, in which loss-of-function mutations of the *GOLIATH* gene are significantly lower than the expected odds. Only nine loss-of-function (LoF) mutations in *GOLIATH* have been described in the gnomAD database, which is 46% of the expected rate of LoF mutation (45). Additionally, there are currently no homozygotes identified for these mutations. Whilst a 70% reduction in *Goliath* mRNA expression in *Goliath*^*–/–*^ mice was sufficient to significantly decrease LDL-C by 45%, the same was not true in heterozygous mice. *Goliath*^*+/–*^ mice had only a 30% reduction in *Goliath* expression, significantly less than the expected 50% reduction, but no significant change in plasma LDL-C. Finally, ASO silencing of *Goliath*, which induced >90% reduction in *Goliath* mRNA levels, elicited the largest decrease in plasma LDL-C levels. Taken together, these data suggest that *Goliath* regulates cholesterol levels in a dose-dependent manner.

In conclusion, our data highlight the complex nature of the post-translational regulation of LDLR. We have identified GOLIATH as a novel post-translational regulator of hepatic LDLR abundance and activity, that may have important implications for targeting the LDLR pathway to lower plasma LDL-C levels. We further provide pre-clinical evidence that the GOLIATH-LDLR pathway could be targeted to enhance LDL clearance from the circulation.

## Methods

### Mice, diets, and treatments

Wildtype mice were purchased from The Jackson Laboratory (JAX, #00664). *Ldlr*^*–/–*^ and *Pcsk9*^*–/–*^ mice on a C57BL/6 background were originally purchased from The Jackson Laboratory. Mice with a targeted disruption of *Goliath* were generated using embryonic stem cells from the European Conditional Mouse Mutagenesis program (EUCOMM) and the European Mouse Mutant Cell Repository (EuMMCR) with a knockout first cassette targeting exon 3 of *Goliath* mRNA. To obtain germline transmission of the *Goliath* KO-first allele (*Goliath*^*KO1*^) on a C57BL/6 background, *Goliath*^*KO1*^ ES cells were injected into blastocysts and the resulting chimeras were bred with C57BL/6 mice. Germline transmission was confirmed by PCR genotyping for a minimum of ten litters. All animals were maintained on normal rodent diet (Ralston Purina Company, 5001) at UCLA on a 12-hour/12-hour light/dark cycle with unlimited access to food and water. Human GOLIATH was cloned as described below and shuttled into adenoviral expression vectors. Adenovirus particles were prepared using the AdEasy system (Agilent) and purified by Cesium Chloride (CsCl) gradient centrifugation. The virus was dialyzed for 48 hours and stored at −80°C. Particles were quantified by serial dilution methods by detection of plaques in HEK293Ad (Agilent) cells. Adenovirus (10^9^ FPU) was delivered by tail-vein injection. The plasmids required for the manufacture of adeno-associated virus (AAV), pAdDeltaF6 adeno helper plasmid (PL-F-PVADF6) and pAAV2/8 (PL-T-PV0007), were obtained from the University of Pennsylvania Vector Core. AAV was prepared by the triple-transfection method (46) in HEK293T cells (ATCC, CRL-3216) and purified by CsCl gradient as previously described (47). Aliquots of concentrated virus were stored at −80°C until injection. Viral titers were determined by qPCR following DNase digestion and normalized to a standard curve of known genome copies. AAV (5 × 10^11^ genome copies) was delivered by intraperitoneal injection. A guide RNA sequence specific to *Goliath* was used to target mouse *Goliath* by CRISPR/Cas9 (5’gactgcacagtgatcgaagt). Gen. 2.5 16-mer antisense oligonucleotides (ASO) targeted to mouse *Goliath* (5’attctgttatcatgac) or control sequences were synthesized and purified by Ionis Pharmaceuticals (Carlsbad, CA) as previously described. ASOs were delivered by intraperitoneal injection twice weekly at a dose of 25 mg/kg for 4 weeks (as described in the figure legends).

### Analysis of CRISPR-based gene disruption

Liver genomic DNA was isolated using the Qiagen DNeasy kit. DNA (100ng) was subjected to qPCR using primers specific to the AAV-Control and AAV-gRNA vectors to assess the relative amount of virus present in the liver. A standard curve from plasmids used for virus production was used to quantify AAV genomes per μg of DNA.

Off-target sites for *Goliath* guide RNA were determined using the online bioinformatics tool, COSMID at https://crispr.bme.gatech.edu/ as previously described (47). Briefly. For on-target editing, 50 μg cDNA was amplified using primers surrounding the gRNA target site. The resulting band was gel-purified (Clontech) and sequenced by Sanger Sequencing. Editing efficiency was estimated using Synthego ICE (Inference of CRIPSR Edits; https://www.biorxiv.org/content/early/2019/01/14/251082).

### Plasma analysis

HDL and LDL/VLDL fractions were separated using a manganese chloride-heparin precipitation (48). Subsequent fractions and total cholesterol were measured using a colorimetric assay (Infinity Cholesterol Reagent; Thermo Scientific). Plasma cholesterol lipoprotein profiles were determined from individual plasma samples using the modified CLiP method as previously described (49, 50).

### RNA analysis

Liver tissue was homogenized, and total RNA extracted using QIAzol reagent (Invitrogen Life Technologies). 500ng total RNA was reverse transcribed using the High Capacity cDNA Reverse Transcriptase Kit (Applied Biosystems) and gene expression determined using a Lightcycler480 Real-time qPCR machine and SYBR-Green mastermix (Roche). Relative gene expression was determined using an efficiency corrected method and efficiency was determined from a 3-log serial dilutions standard curve made from cDNA pooled from all samples. Results were normalized to *36b4* mRNA. Primer sequences are available upon request.

### Protein analysis

Liver tissue was homogenized, and protein extracted using RIPA buffer with protease inhibitor cocktail mix (1 Complete MINI EDTA-free protease inhibitor tablet (Roche), 25 μg/mL calpain inhibitor (Sigma), and 200 μM PMSF (Sigma)). 25-50 μg protein was separated on an SDS-PAGE gel (BioRad) and transferred to a polyvinylidene fluoride membrane. Membranes were incubated overnight with antibodies to GOLIATH (1:1000, sigma), EGFR (1:1,000; Invitrogen), FLAG (1:1,000; Sigma), GFP (1:1,000; Santa Cruz Biotechnology), HA.11 (1:1,000; Covance), LDLR (1:1,000; Cayman), LRP1 (1:1,000; Abcam), PDI (1:1,000; Cell Signaling), and Transferrin Receptor (1:1,000; Invitrogen). Proteins were detected with HRP-conjugated secondary antibodies (1:10,000; GE Healthcare) and visualized using an AI600 Imager (GE Healthcare). Densitometry analysis was performed using ImageQuant (TL 8.1, GE Healthcare).

### Plasmids and expression constructs

The pDEST47-hLDLR and K1/6/20RC29A mutant plasmids were a kind gift from Dr. Peter Tontonoz (37, 51). The pcDNA3.1-(HA-Ubiquitin)_6_ plasmid was a kind gift from Dr. James Wohlschlegel (UCLA, USA). Human GOLIATH was amplified (Fwd 5’ atgagctgcgcggggcgggcgggccctgccc; Rev 5’ gcttgaatgctaatgaggtagaatggttttga) from Hep3B cell cDNA using HiFi DNA polymerase (KAPA Biosciences). Human GOLIATH was also generated with a 3xFLAG epitope engineered at amino acid position 28 after the signal peptide as a double-stranded DNA gBlock (IDT Technologies). Additionally, the C304A mutation was introduced into the RING domain of GOLIATH. Restriction digests and DNA sequencing were used to confirm all of the constructs used in this study. For adenoviral studies, full length human GOLIATH was sub-cloned from pcDNA3.1 into the pAd-CMV vector.

### Guide RNA Design

*Staphylococcus aureus* guide RNAs (gRNAs) targeting murine *Goliath* were designed by examining the coding sequence using SnapGene. Preliminary off target prediction was completed using CRISPR Off-target Sites with Mismatches, Insertions and/or Deletions (COSMID) (52). The search terms used for the off-target search aimed to return the highest possible number of off-targets, using a NNGRR protospacer adjacent motif (PAM) instead of NNGRRT, and allowing three mismatches and two insertions or deletions within the gRNA and PAM sequence. Of the available gRNAs, we selected, cloned, and tested the gRNAs with the fewest predicted off-targets. The plasmid 1313 pAAV-U6-BbsI-MluI-gRNA-SA-HLP-SACas9-HA-OLLAS-spA (Addgene #109304) was a gift from Dr. William Lagor at Baylor College of Medicine (53). This AAV vector backbone contains inverted terminal repeats surrounding a U6 promoter driving gRNA expression and a hybrid liver-specific promoter (HLP) driving expression of *Staphylococcus aureus* Cas9 (SaCas9). The gRNA was cloned into this vector by ligating annealed oligos matching the desired gRNA sequence into the BbsI cloning site behind the U6 promoter. The Control AAV-CRISPR used in experiments is the parent vector containing the BbsI cloning site. When the sequence including the BbsI site is queried using COSMID for the SaCas9 NNGRR PAM, no possible off-targets are returned.

### Cell culture and transfection

HEK293T cells were obtained from ATCC (CRL-3216). HEK293T cells were maintained in DMEM containing 10% FBS. For transfections, cells were plated onto cell-culture-treated plates or coverslips (Corning) at 60% confluence on Day 0. Cells were transfected with a total of 1μg combined DNA plasmids using FuGENEHD (Promega) according to the manufacturer’s instructions. After 24-48hr incubation, cells were collected for downstream processing (western blot, immunoprecipitation) or fixed and mounted for imaging. Quantification of confocal images was performed using ImageJ.

### Cell Surface Biotinylation

For cell surface biotinylation, HEK293T cells were transfected with plasmids encoding GFP-tagged LDLR, HA-tagged ubiquitin, and N-terminally FLAG-tagged WT or mutant (C304A) GOLIATH as described above. After 36 hours, cells were washed in PBS++ (Phosphate buffered saline with 0.02mM CaCl_2_ and 0.15mM MgCl_2_) and then incubated for 30 minutes on ice with EZ-link SulfoNHS-SS Biotin (diluted in PBS++). The cells were washed in PBS++ and the reaction was quenched for 30 minutes at 4°C in quenching buffer (PBS++ with 100mM glycine). Biotin-modified proteins were immunoprecipitated with NeutrAvidin streptavidin beads overnight at 4°C. The following day, biotin-modified proteins were collected by centrifugation at 5,000 × g for 5 min at 4°C. Intracellular, unmodified proteins were collected from the supernatant of the 5,000 × g spin. The streptavidin beads were washed three times in PBS++ before proteins were removed from the beads by incubation at 42°C for 20 min in Laemmli sample loading buffer supplemented with β-mercaptoethanol. Equal % volume of individual fractions were subject to immunoblotting.

### Immunoprecipitation

Cells were plated and transfected as described above. Total cell lysates were prepared in RIPA buffer as described above. Lysates were cleared by centrifugation at 4°C for 10 min at 10,000 x *g*. Protein concentration was determined using the BCA Assay (BioRad) with bovine serum albumin as a reference. To immunoprecipitate LDLR-GFP, equal amounts of protein of cleared lysate were incubated with anti-GFP polysera (1:1,000) overnight prior to addition of protein-G agarose beads (Santa Cruz Biotechnology) for an additional 2 hr. Subsequently, beads were washed 3x with RIPA buffer supplemented with protease inhibitors. All incubations and washes were done at 4°C with rotation. Proteins were eluted from the beads by incubating in Laemmli sample buffer at 70°C for 30 min.

### Imaging

Cells were plated and transfected as described above. At harvest, cells were washed, fixed in 4% PFA for 15 min, and then washed again before briefly being stored in PBS. Coverslips were mounted on to slides imaged using a 63X lens under immersion oil. Images were captured using a Zeiss AxioCam 506 camera and post-image analysis and quantification was carried out using ImageJ.

### Statistical Analysis

Statistical analysis was performed using Prism Graphpad software (V8.0). All results are mean ± SD or SEM, as stated in the figure legends. Normality was determined using D’Agostino-Pearson and/or Shapiro-Wilk normality testing. *P* values for normally distributed data were calculated using either a two-tailed Student’s *t*-test, or one-way or two-way ANOVA with either Tukey’s or Sidak’s post hoc analysis as indicated in the figure legends. *P* values for non-normally distributed data were calculated using the Mann-Whitney rank sum test of the Kruskal-Wallis test with Dunn’s multiple comparisons testing. A *P* < 0.05 was considered significant and statistical significance is shown as described in the figure legends.

### Study Approval

All animal experiments were approved by the Office of Animal Research Oversight (OARO) and the Institutional Animal Care and Use Committee (IACUC) at the University of California Los Angeles.

## Author Contributions

E.J.T. and T.Q. de A.V. conceived and oversaw the project. B.L.C., K.E.J., J.C., A.C., M.S., P.M., T.Q. de A.V., and E.J.T performed experiments. K.E.J. designed and purified AAV-CRISPR. B.L.C. and K.E.J. designed and performed AAV-CRISPR experiments. J.C. and A.C. assisted with animal experiments. P.M. assisted with western blotting. M.S. performed functional genomic analyses. A.B. performed FPLC lipoprotein profiles. R.L. provided antisense oligonucleotide compounds. Data analysis and statistical analyses were performed by B.L.C. and E.J.T. Figures were generated by B.L.C. and E.J.T. The manuscript was written by B.L.C. and E.J.T. All authors revised and approved the final manuscript.

## Declaration of Interests

The authors declare no conflict of interest.

## Acknowledgements

We thank Rachel Scott and David Merriott for help with *in vitro* studies. We thank Ionis Pharmaceuticals for providing antisense oligonucleotide compounds. We thank all members of the Tarling-Vallim and Tontonoz labs at UCLA for useful advice, discussion, and sharing reagents. We thank Peter Edwards for discussion and review of the manuscript. B.L.C. is sponsored by an AHA post-doctoral fellowship (19POST34380145). K.E.J. is sponsored by the UCLA Vascular Biology T32 fellowship (5T32HL069766). M.S. is funded by NIH grant HL138193. A.B. is funded by NIH grants DK125048 and HL107794. T.Q. de A.V. is funded by NIH grants HL122677, DK 119112, and DK118064. E.J.T. is funded by NIH grant HL136543.

## Notes

### Competing Interest Statement

The authors have declared no competing interest.

