## Supplemental Figures for "GOLIATH regulates LDLR availability and plasma LDL cholesterol levels"

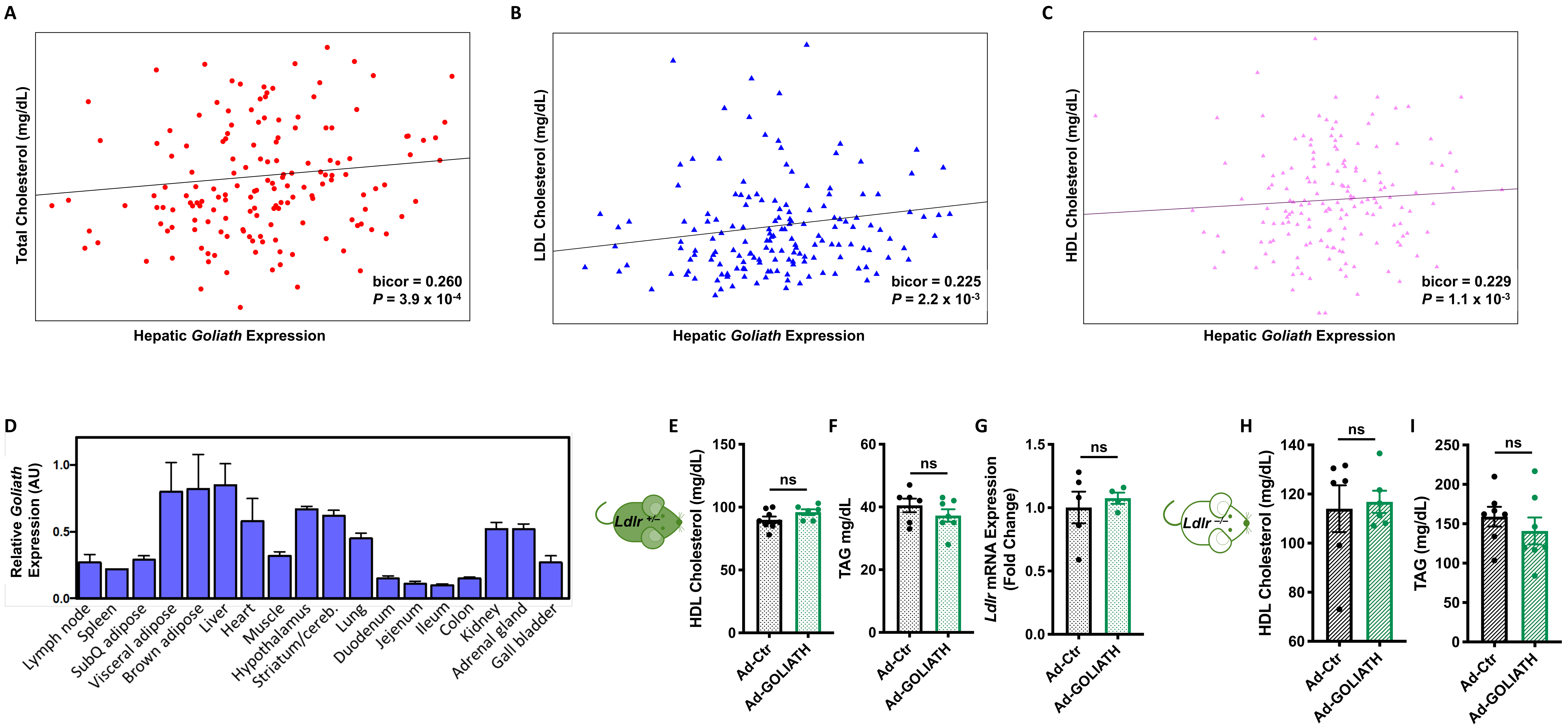


**Clifford et al. Supplemental Figure 1. GOLIATH reduces plasma LDL-C levels *in vivo* in a process that is dependent on LDLR.** Mouse hepatic *Goliath* mRNA expression is significantly correlated with plasma **(A)** total cholesterol, **(B)** LDL-cholesterol, and **(C)** HDL-cholesterol levels. **(D)** *Goliath* mRNA levels in mouse tissues (n=3 mice/tissue). **(E, H)** Plasma HDL cholesterol in *Ldlr*$^{+/-}$ mice **(E)** and *Ldlr*$^{-/-}$ mice **(H)** treated with Ad-Ctr or Ad-GOLIATH as in Figure 1 (*n*=5-8 mice/group). **(F, I)** Plasma triglyceride (TAG) in *Ldlr*$^{+/-}$ mice **(F)** and *Ldlr*$^{-/-}$ mice **(I)** treated with Ad-Ctr or Ad-GOLIATH as in Figure 1 (*n*=5-8 mice/group). **(G)** Hepatic *Ldlr* mRNA expression in *Ldlr*$^{+/-}$ mice treated with Ad-Ctr or Ad-GOLIATH as in Figure 1 (*n*=5-8 mice/group). Data are expressed as mean ±SEM with individual animals noted as dots. Statistical significance was determined by Student's *t* test. ns, not significant.

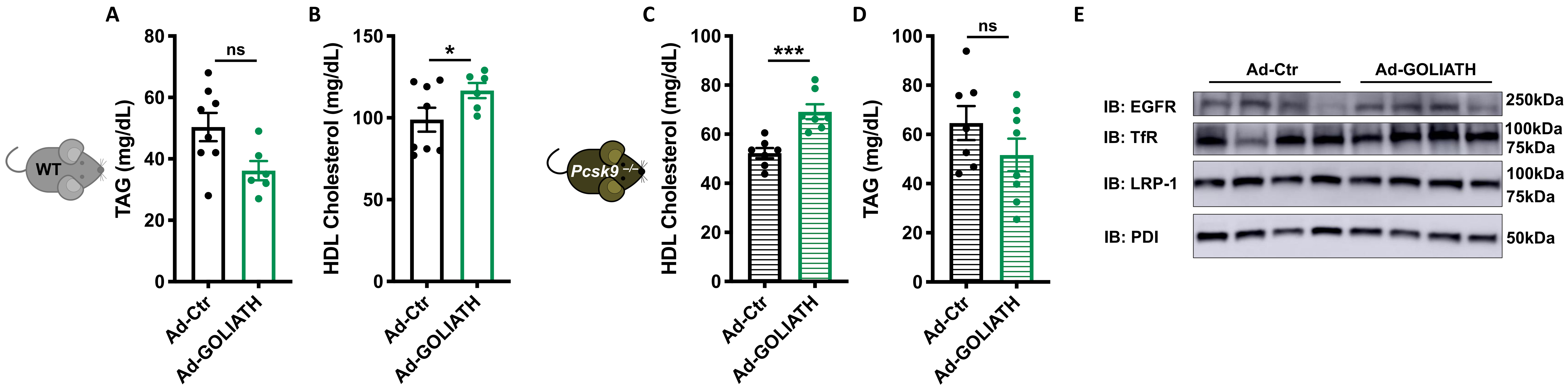


**Clifford et al. Supplemental Figure 2. GOLIATH regulates plasma cholesterol levels in wildtype and *Ldlr*$^{+/–}$ mice.**
Wildtype (WT) **(A-B)** and *Pcsk9*$^{–/–}$ **(C-D)** mice were treated with control adenovirus (Ad-Ctr) or adenovirus overexpressing human GOLIATH (Ad-GOLIATH) for 7 days (*n*=5-8 mice/group) as in Figure 2. Plasma HDL cholesterol **(A)** and triglycerides (TAG) **(B)** in wildtype mice treated with Ad-Ctr or Ad-GOLIATH. Plasma HDL cholesterol **(C)** and TAG **(D)** in *Pcsk9*$^{–/–}$ mice treated with Ad-Ctr or Ad-GOLIATH. **(E)** Western blots of total liver extracts using antibodies to LRP1, EGFR, and Transferrin Receptor. Data are expressed as mean ± SEM with individual animals noted as dots. Statistical significance was determined by Student's *t* test. * $p<0.05$, *** $p<0.001$. ns, not significant. EGFR; Epidermal Growth Factor Receptor; LRP1, LDLR-related protein 1; TfR, Transferrin Receptor; PDI, protein disulfide isomerase.

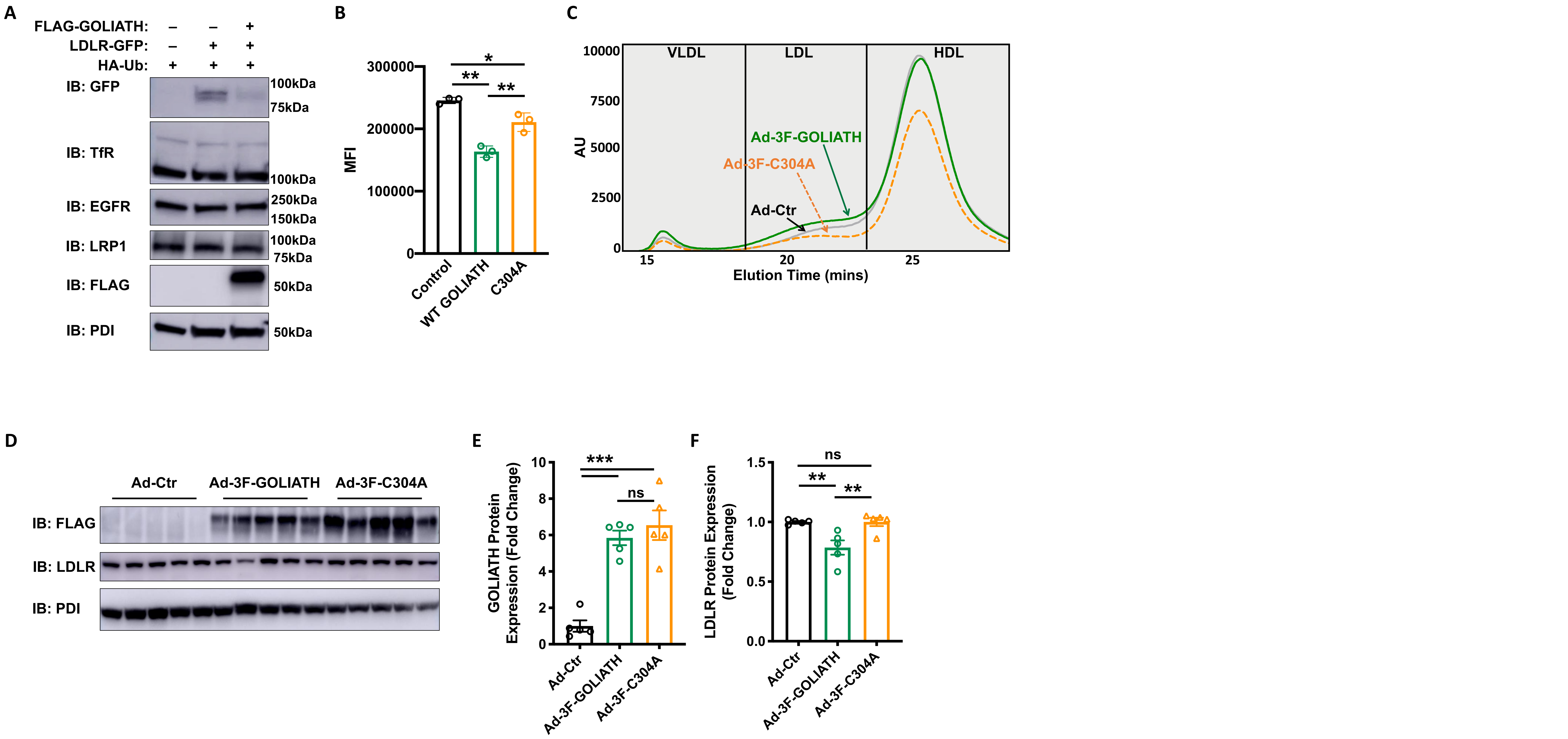


**Clifford et al. Supplemental Figure 3. Regulation of plasma cholesterol is dependent on GOLIATH E3 ubiquitin ligase function.**
**(A)** HEK293 cells were co-transfected with LDLR-GFP and FLAG-GOLIATH. After 36 hours, lysates were subjected to IB. EGFR, epidermal growth factor receptor; TfR, transferrin receptor, LRP1, LDLR-related protein 1, PDI, protein disulfide isomerase. **(B)** Mean fluorescence intensity quantification of confocal images in Figure 3E. n=3 images from 3 independent transfections. **(C)** FPLC lipoprotein profiles of wildtype mice treated as in Figure 3F. **(D-F)** Total lysates from liver from mice treated as in Figure 3F were analyzed by immunoblotting (IB). **(D)** and protein expression of **(E)** 3xFLAG-GOLIATH and **(F)** LDLR was quantified by densitometry. Data are expressed as mean ±SEM with individual animals noted as dots. Statistical significance was determined by one-way ANOVA. *$p < 0.05$, ** $p < 0.01$, *** $p < 0.001$, ns, not significant.

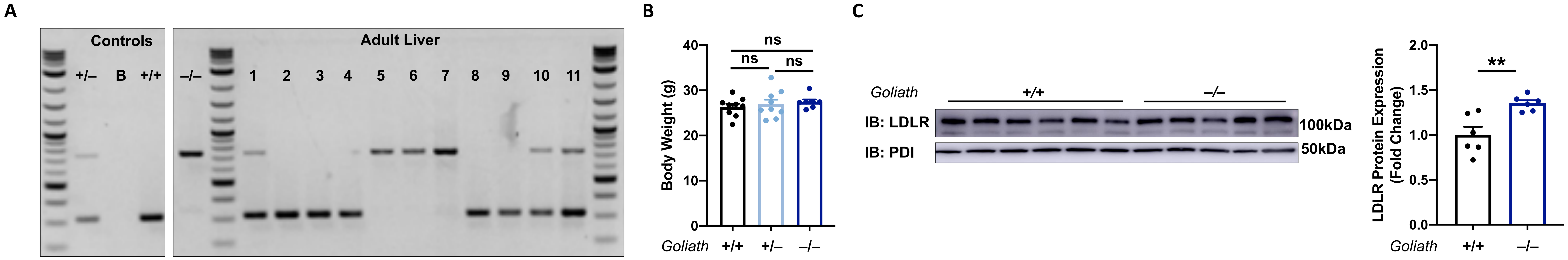


**Clifford et al. Supplemental Figure 4. Generation of *Goliath* KO mice.**

**(A)** Genotyping for adult liver from *Goliath*$^{+/+}$, *Goliath*$^{+/–}$, and *Goliath*$^{–/–}$ mice. B, Blank. **(B)** Body weight of 10-week old *Goliath*$^{+/+}$, *Goliath*$^{+/–}$, and *Goliath*$^{–/–}$ mice. **(C)** Representative western blot and quantification of hepatic protein expression of LDLR in *Goliath*$^{+/+}$ and *Goliath*$^{–/–}$ mice. Data are expressed as mean ±SEM with individual animals represented as dots. (n=5-10 mice/genotype). Significance was measured by one-way ANOVA. ** $p < 0.01$. ns, not significant.

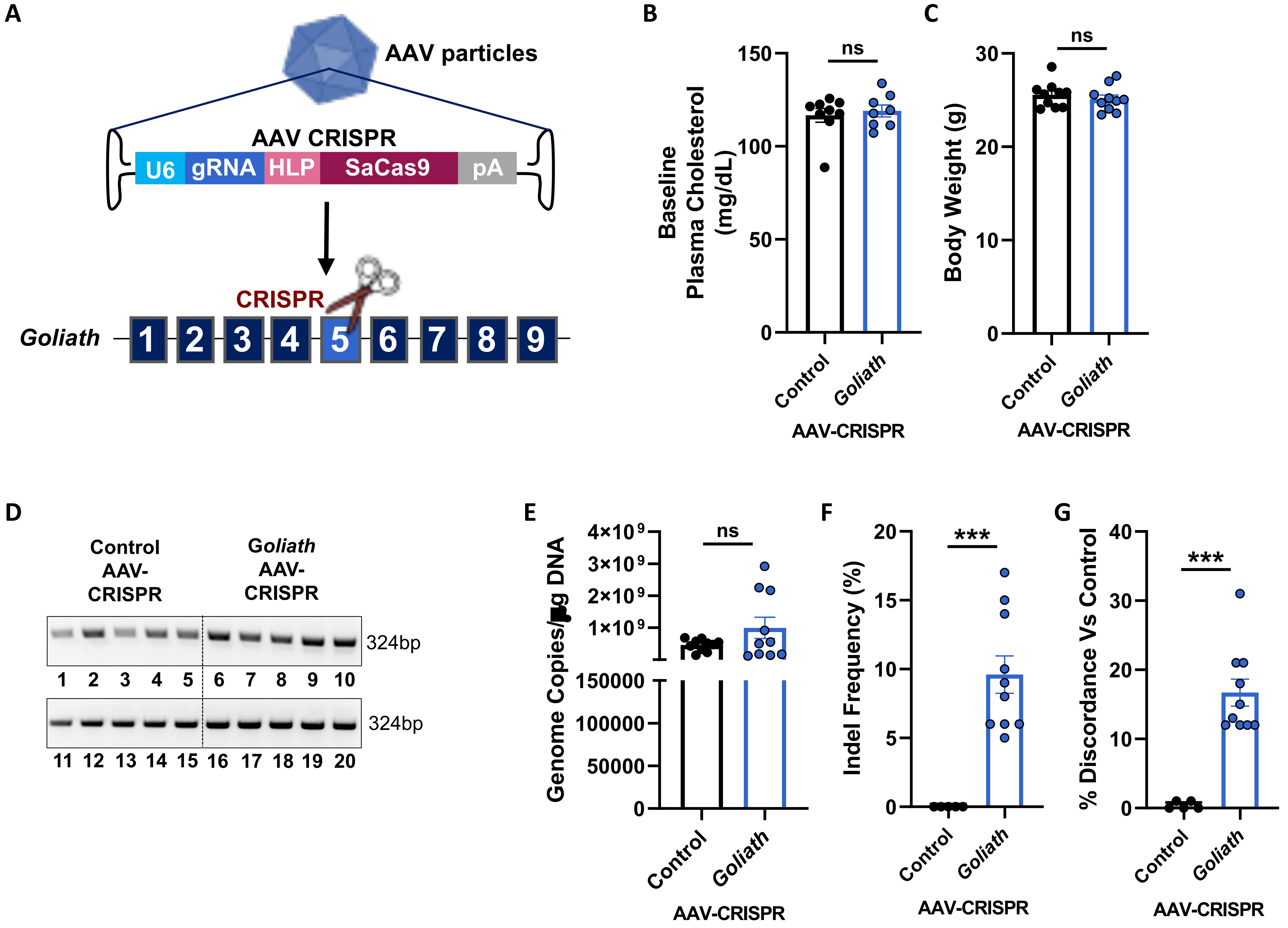


**Clifford et al. Supplemental Figure 5. Liver-specific disruption of *Goliath* reduces plasma cholesterol levels.**
**(A)** Schematic of the CRISPR disruption strategy. An all-in-one AAV-CRISPR was used to express a gRNA targeting exon 5 of *Goliath* and SaCas9. **(B)** Baseline plasma cholesterol and **(C)** Body weight of wildtype mice treated as in Figure 5. **(D)** Confirmation of viral load in the livers of mice as in Figure 5 determined by PCR amplifying within hSaCas9. **(E)** Vector genomes present in the livers of mice injected with Control AAV-CRISPR or *Goliath* AAV-CRISPR from Figure 5 as determined by qPCR. Editing analyses were completed using Synthego ICE on mRNA from Control AAV-CRISPR and *Goliath* AAV-CRISPR treated animals with **(F)** ICE INDEL efficiency and **(G)** ICE sequence discordance demonstrating on-target editing still present in ~10-20% of the remaining mRNA**.** The remaining 70% of the mRNA were likely degraded. Data are expressed as mean ±SEM with individual animals noted as dots. Significance was measured by Student's *t*-test. ns, not significant.
